# Comparative study of the mechanism of natural compounds with similar structures using docking and transcriptome data for improving in silico herbal medicine experimentations

**DOI:** 10.1101/2023.04.23.538005

**Authors:** Musun Park, Su-Jin Baek, Sang-Min Park, Jin-Mu Yi, Seongwon Cha

**Affiliations:** Korean Medicine (KM) Data Division, Korea Institute of Oriental Medicine, Daejeon, Republic of Korea; College of Pharmacy, Chungnam National University, Daejeon, Republic of Korea; KM Convergence Research Division, Korea Institute of Oriental Medicine, Daejeon, Republic of Korea

**Keywords:** natural compounds, herbal medicine, similar structure analysis, molecular docking, transcriptome

## Abstract

Natural products have successfully treated several diseases using a multi-component, multi-target mechanism. However, a precise mechanism of action has not been identified. Systems pharmacology methods have been used to overcome these challenges. However, there is a limitation as those similar mechanisms of similar components cannot be identified. In this study, comparisons of physicochemical descriptors, large-scale molecular docking analysis, and RNA-seq analysis were performed to compare the mechanisms of action of similar compounds and to confirm the changes observed when similar compounds were mixed and used. We propose an advanced method for in silico experiments in herbal medicine research based on the results. First, physicochemical descriptors were calculated based on the chemical structures of oleanolic acid (OA), hederagenin (HG), and gallic acid (GA). Similarities were confirmed by calculating the Euclidean, cosine, and Tanimoto distances between the descriptors. Next, the mechanisms of action of OA, HG, and GA were compared and confirmed through in silico-based systems pharmacology analysis using the BATMAN-TCM platform. The proteins interacting with the three compounds were verified through large-scale molecular docking analysis using the druggable proteome. Finally, a drug response transcriptome study was performed using OA, HG, GA, and a combination of OA and HG (COH) with similar structures.A comparison of physicochemical descriptors confirmed that OA and HG were very close. In particular, the two compounds showed a concordance rate of > 99% at cosine and Tanimoto distances. The systems pharmacology analysis results confirmed that OA and HG shared more than 86% of their predicted target proteins and differed only in GA. Systems pharmacology analysis revealed that OA and HG share the mechanisms of cardiac muscle contraction, oxidative phosphorylation, and non-alcoholic fatty liver disease. In a molecular docking analysis of the 50 major druggable proteins, OA and HG shared 38 proteins, while GA shared a few with proteins derived from the other two compounds. In addition, OA and HG were confirmed to act on gonadotropin-releasing hormone (GnRH) secretion, type 2 diabetes mellitus, cholinergic synapses, and calcium signaling pathways, and docking analysis visualization confirmed that the two components interact at the same site. RNA-seq analysis also showed that the differentially expressed genes and pathways derived from OA and HG were similar, and it was confirmed that COH had similar results to OA and HG. Our study has three novel findings. First, an advanced network pharmacology research method was suggested by partially presenting a solution to the difficulty in identifying multicomponent mechanisms. Second, a new natural product analysis method was proposed using large-scale molecular docking analysis. Finally, various biological data and analysis methods were used, such as in silico system pharmacology, docking analysis, and drug response RNA-seq. The results of this study are meaningful in that they suggest an analysis strategy that can improve existing systems pharmacology research analysis methods by showing that natural product-derived compounds with the same scaffold have the same mechanism.

## 1. Introduction

Natural products have long been used to treat patients in Traditional Asian, Ayurvedic, and Kampo medicine. Therefore, it can be said that natural products are safe and effective (1). However, the exact mechanism underlying the therapeutic efficacy of natural compounds has not yet been identified, which is an obstacle to the commercialization of natural product-based drugs. (2). If this issue is not overcome soon, it will not only result in the loss of trust in treatments using natural products but will also hinder the standardization and industrialization of natural products. Therefore, it is important to understand the mechanisms of action of these natural products.

Most mechanisms of action studies have been conducted as single compound-single target studies based on the Magic Bullet Paradigm (3, 4). However, because natural products contain multiple compounds, studies that consider only a single compound have limitations in revealing the mechanisms of action of natural products. To overcome this limitation, network-based systems pharmacology studies are being conducted to verify the mechanisms of action of natural products based on multi-component, multi-target interactions (5, 6, 7). Although herbal medicine research using systems pharmacology has been successfully conducted on platforms such as TCMSP (8) and BATMAN-TCM (9), it still has the limitation of showing interaction results without considering similar compounds.

Active compounds with similar structures have similar mechanisms of action. For example, caffeine has the same graph framework as adenosine and exerts competitive antagonistic effects on adenosine receptors (9, 10). In addition, based on the finding that the same molecular scaffold has the same mechanism of action (11, 12), research on the development of new drugs using fragment-based drug design, such as scaffold hopping, has also been conducted (13, 14, 15). These studies show that comparative studies on similar compounds are necessary to verify the mechanism of action of natural products containing many similar active compounds.

Natural products contain several compounds with similar structures. These include compounds such as peptides, polyketides, and terpenes. Terpenes are phytochemicals with various physiological activities (16) and exist in various forms in natural products through structural diversification by cytochrome P450 enzymes (CYP) (17). For example, oleanolic acid, a pentacyclic triterpene present in *Paeonia lactiflora*, is biologically transformed by CYP into compounds such as hederagenin, gypsogenic acid, and medicagenic acid (17). However, their structures are maintained because the biological transformation processes do not change the molecular scaffold but the functional group. This suggests that, although natural products contain various compounds owing to the biotransformation process, the transformed components have the same scaffold and mechanism of action.

One analytical method for verifying this hypothesis is molecular docking. Molecular docking is an in silico method that determines the interaction potential by calculating the binding affinity between a specific protein and a specific compound (18). When compounds with similar structures interact at the same location on a protein, they are likely to share the same biological mechanisms. Therefore, determining whether similar compounds dock to the same protein and match the docking binding site is necessary for drug mechanism studies. In particular, because natural products contain many compounds with similar structures that interact with multiple targets, large-scale molecular docking analysis is important for studying the mechanisms of action of natural products. However, few studies have elucidated the mechanisms of action of these natural products using large-scale molecular docking analyses.

Although large-scale molecular docking analysis succeeds in predicting whether natural products with similar structures have similar mechanisms, it remains unclear whether they have the same mechanisms. In addition, it is necessary to predict the results of drug combinations to reveal the mechanism of action of natural products in which multiple compounds are combined; however, it is difficult to predict the mechanism of drug combinations with similar structures using conventional network pharmacology or docking analysis. Therefore, we attempted to resolve these issues using drug response transcriptome analysis. Transcriptome analysis using RNA-Seq reveals the RNA expression level in cells using next-generation sequencing analysis and can identify intracellular transcripts that change in response to specific stimuli (19). Because the expression of transcripts induced by drug treatment reflects a wide range of target changes, it provides abundant information on drug action mechanisms (20). Using the characteristics of these drug response transcripts, it is possible to identify the mechanism of action of a combination of natural compounds with similar structures.

While researching the mechanisms of natural products, it is important to consider the similarity of compound structures within natural products; however, few studies have analyzed and compared natural product compounds with similar structures. Therefore, predicting the biological mechanisms of compounds with similar structures and identifying the drug mechanisms of similar compound combinations is novel. This study proposes an advanced method for in silico experiments for herbal medicine research. The method involves comparing the mechanisms of action of natural compounds with similar structures using biological data.

## 2. Methods

### 2.1 Overview of the study

This study was conducted in the following manner: Physicochemical descriptors were calculated based on the chemical structures of oleanolic acid (OA), hederagenin (HG), and gallic acid (GA), and similarity was confirmed by calculating the distance between the descriptors. Next, the mechanisms of action of OA, HG, and GA were compared and confirmed through pharmacological analyses using in-silico-based systems. In addition, the proteins interacting with the three compounds were verified through large-scale molecular docking analysis using the druggable proteome. Finally, it was confirmed through the production and analysis of drug response transcripts that the mechanism of action of OA and HG was similar and consistent with the combination of OA and HG (Figure 1).

**Figure 1.**
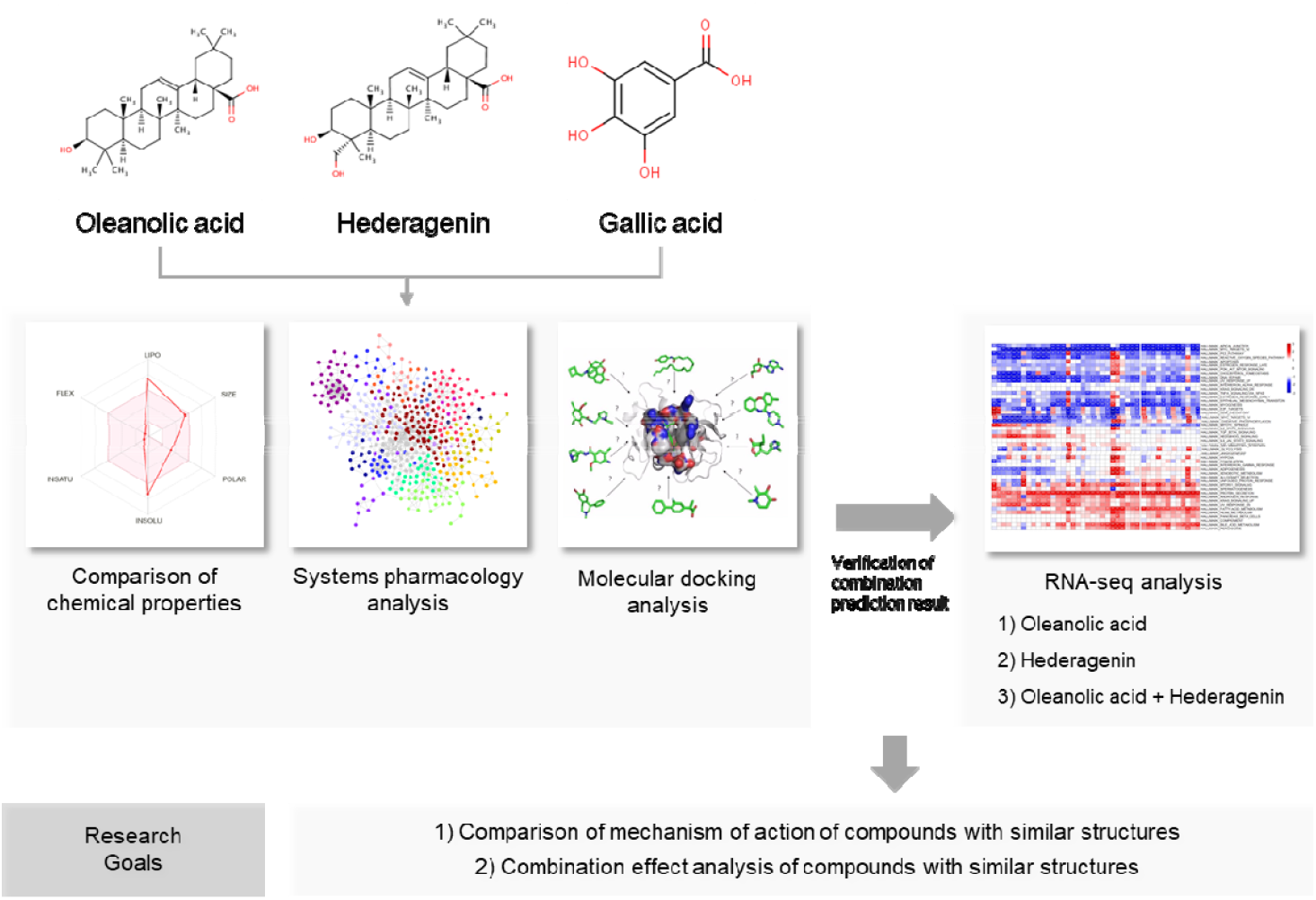
Framework for Bioinformatics Analysis of Structurally Similar Compounds. Research comparing the mechanism of action of natural product compounds with similar structures using biological data and research on the mechanism of action using a combination of natural product components with similar structures was conducted. Oleanolic acid (OA) and Hederagenin (HG) were selected as natural compounds with similar structures, and gallic acid (GA) was selected as a control. To compare the mechanisms of action of OA, HG, and GA, chemical properties comparison, system pharmacology analysis, and molecular docking analysis were performed, respectively. Then, RNA-seq analysis was performed to confirm the mechanism of action of natural compounds with similar structures and the mechanism of action of combinations of natural compounds with similar structures. This study confirmed that compounds of natural products with similar structures have the same mechanism, and even when compounds with similar structures were mixed, it was confirmed that they had the same mechanism.

### 2.2 Comparative Analysis of Chemical Properties of OA, HG, and GA

#### 2.2.1 Molecular Feature Collection and Molecular Descriptor Calculation Method

The physical properties, CID, and SMILES string information of OA, HG, and GA used in this study were collected from the PubChem database, and the chemical ontology information was collected from the ChEBI database (21). Molecular descriptors were calculated using the Python Mordred library (22). Among the 1826 molecular descriptors provided by Mordred, 396 descriptors that could not be calculated, and 314 descriptors with a value of zero for all three compounds were excluded from the analysis. The 1116 molecular descriptors used in the similarity analysis are presented in Supplementary Material 1.

#### 2.2.2 Distance-based Molecular Similarity Measurement Method

The similarity measures for OA, HG, and GA were calculated based on distance measures. The similarity distance measure used 1116 molecular descriptors computed from the Mordred library. Compounds were paired for distance calculation, and the Euclidean [1], cosine [2], and Tanimoto distances [3] of the paired compounds were calculated using the Python numpy library (23).

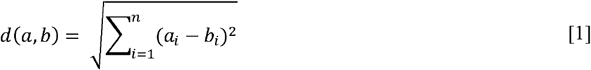

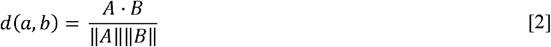

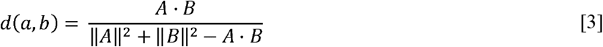

### 2.3 Comparative Analysis of OA, HG, and GA based on Systems Pharmacology Platform

#### 2.3.1 Selection of Druggable Target Using Systems Pharmacology Platform

In the systems pharmacology analysis, protein targets interacting with OA, HG, and GA were selected using the BATMAN-TCM platform. On the BATMAN-TCM platform, the possibility of drug-target interaction (DTI) was calculated and expressed as a score; among them, targets with a DTI score of 10 points and above were selected as druggable targets.

#### 2.3.2 Network Configuration Using Druggable Targets

The druggable targets of each compound were constructed as a compound-target-pathway network. The network nodes consisted of compounds (OA, HG, and GA), targets of effective pathways. The edges of the network represent 1) the connectivity of the compound that interacts with the targets and 2) the connectivity of the target and the pathways containing the target. The target and pathway nodes used in the network were selected using over-representation analysis (ORA) based on the gene sets of the KEGG pathway database (24) provided by the BATMAN-TCM database. The effective pathways for each compound were determined based on an adjusted p-value less than 0.05, calculated using the Benjamini-Hochberg (BH) procedure (25). Along with the effective pathways, druggable targets belonging to effective pathways were selected as network nodes. Network visualization was performed using Cytoscape (v3.9.1) (26).

#### 2.3.3 Over-representation Analysis Based on Systems Pharmacology Platform

Druggable targets OA, HG, and GA were used for ORA. ORA was performed on the EnrichR platform (27) and analyzed using gene sets of the KEGG pathway (24), the gene sets of GO biological process (28), and the gene sets of OMIM disease (29). The analysis results were based on the combined score provided by the EnrichR platform, and the top 10 pathways and diseases were selected as the pharmacological mechanisms of each compound.

### 2.4 Molecular Docking Analysis

#### 2.4.1 Molecular Docking Analysis Method

The compounds used for docking analysis were transformed into pdbqt form using OpenBabel software (30) after downloading the 3d_sdf information from the PubChem database (31) (PubChem CID: OA-10494, HG-73299, and GA-370). The protein structures used for analysis were collected from the AlphaFold (AF) (32) and Human Protein Atlas databases (HPA) (33). Among the human-derived proteomes provided by the AF, 812 proteomes selected by the HPA as druggable proteomes were used for analysis. After converting the selected druggable proteomes to pdbqt, which can be docked using the OpenBabel Python API, molecular docking analysis was performed with the modified compound. Molecular docking analysis was performed using the Python and AutoDock Vina software (32), and the docking analysis parameter exhaustiveness was set to a maximum value of 100.

#### 2.4.2 Pathway Analysis and Visualization Using Molecular Docking Analysis Results

The druggable proteomes of OA, HG, and GA, selected through docking analysis, were used for ORA-based pathway analysis using the EnrichR platform. Fifty proteins with the lowest binding affinities were selected as docking-based effective proteins (DEPs), and ORA was performed. Similar to the systems pharmacology platform method, ORA was performed using gene sets from the KEGG pathway, GO Biological Process, and OMIM Disease databases. The top 10 pathways and diseases with the highest combined scores were selected as the DEP-based pharmacological mechanisms of each compound.

Pathway visualization was performed by constructing a network using Cytoscape software. The five pathways with the highest combined scores were selected as the main active pathways from the KEGG pathway analysis results calculated using EnrichR for nrtwork construction. The nodes of the network were composed of each compound, DEPs, and the main active pathways. The edges of the network comprised interactions between compounds and DEPs and interactions between DEPs and major active pathways.

The docking analysis was visualized by selecting five proteins (ABL1, JAK1, PIK3CD, CPS1, and CACNA1S) that effectively interacted with all three components in the docking analysis. RYR1 was excluded from the visualization because its sequence length was too large, and it was split into multiple files in the AlphaFold database to provide sequence information. AutoDock tools were used to visualize the molecular docking prediction results for the five proteins and each compound (33).

### 2.5 RNA-seq Analysis Method

#### 2.5.1 Chemicals and Reagents

Roswell Park Memorial Institute (RPMI) 1640 medium, phosphate-buffered saline (PBS), TrypLE Express, penicillin-streptomycin, and fetal bovine serum (FBS) were purchased from Gibco (Grand Island, NY, USA). Cell culture flasks and multiwell culture plates were purchased from Thermo Fisher Scientific (Waltham, MA, USA). Dimethyl sulfoxide (DMSO) and QIAzol lysis reagents were purchased from Sigma-Aldrich (St. Louis, MO, USA) and Qiagen (Germantown, MD, USA), respectively. The Ez-Cytox cell viability assay kit was purchased from Dogen Bio (Seoul, Korea). Oleanolic acid (CFN98800), hederagenin (CFN98695), and gallic acid (CFN99624) were purchased from ChemFace (Wuhan, Hubei, China) and all compounds had purities ≥ 98%.

#### 2.5.2 Cell Culture

Human non-small cell lung cancer cells (NSCLC) line A549 (CCL-185) was purchased from the American Type Culture Collection (ATCC, Manassas, VA, USA). The cells were maintained in RPMI 1640 medium supplemented with 10% (v/v) heat-inactivated FBS, 100 IU/mL penicillin, and 100 mg/mL streptomycin at 37 ° C, 5% CO_2_ incubator. A549 cells were subcultured every 3 or 4 days, depending on the cell density.

#### 2.5.3 Drug Treatment and Total RNA Preparation

The compounds were dissolved at 20 mM in DMSO and stored at –20 °C until use. Before drug treatment, 20 mM drug solutions were diluted to 100 and 200 μM with PBS, and filtered through a 0.22-μm membrane syringe filter (Sartorius, Goettingen, Germany). PBS with 2% DMSO was used as the vehicle. A549 cells were plated at 3 × 10^5^ cells/well in a 6-well plate containing 3 mL growth medium one day before drug treatment. The cells were exposed to 5 or 10 μM by treatment with 150 uL of 100 or 200 μM diluted drug per well. There was no cytotoxicity when treated with a high dose (10 μM) in all drugs, and it was confirmed using the Ez-cytox cell viability assay kit. After 24 h of drug treatment, the cells were washed thrice with ice-cold PBS. The total cell lysate was prepared with QIAzol lysis reagent and stored in a -70 °C deep freezer until RNA extraction. Total RNA was isolated according to the manufacturer’s protocol. The concentration of the isolated RNA was determined using an Agilent RNA 6000 Nano Kit (Agilent Technologies, Waldbronn, Germany), and RNA quality was evaluated by assessing the RNA integrity number (RIN > 7).

#### 2.5.4 Library Preparation for mRNA Sequencing

One milligram of total RNA was processed to prepare an mRNA sequencing library using the MGIEasy RNA Directional Library Prep Kit (MGI) according to the manufacturer’s instructions. The first step involved purifying poly A-containing mRNA molecules using poly T oligo-attached magnetic beads. Following purification, mRNA was fragmented into small pieces using divalent cations at elevated temperatures. Cleaved RNA fragments were copied into first-strand cDNA using reverse transcriptase and random primers. Strand specificity was achieved in RT directional buffer, followed by second-strand cDNA synthesis. A single ‘A’ base was then added to these cDNA fragments, and subsequent ligation of the adapter occurred. The products were purified and enriched using PCR to create a final cDNA library. The double-stranded library was quantified using QauntiFluor ONE dsDNA System (Promega). The library was circularized at 37 °C for 30 min and then digested at 37 °C for 30 min, followed by the cleanup of the circularization products. DNA nanoballs (DNB) were prepared by incubating the library at 30 °C for 25 min with DNB. Finally, the library was quantified using the QuantiFluor ssDNA System (Promega).

#### 2.5.5 Sequencing and Estimate Expression Abundance

The prepared DNB was sequenced using the MGIseq system (MGI) with 100 bp paired-end reads. Reads were trimmed using Trim Galore (34) to remove adapter sequences and low-quality reads. High-quality sequence reads were mapped to the human genome (hg38), and mRNA expression levels were quantified using the DESeq2 library (35).

### 2.6 Gene Set Enrichment Analysis and Cluster Analysis of Pathways

Gene set enrichment analysis (GSEA) was performed using the RNA sequencing results obtained from transcriptome experiments (36). Differentially expressed genes (DEGs) were extracted using DESeq2 library (v1.38.2) (35) and edgeR (v3.40.1) (37) included in Bioconductor (38), and the extracted DEGs were visualized using volcano plots (DEGs selection threshold: q-value ≤ 0.05, p-value ≤ log2(1.5)). GSEA was performed for curated gene sets (KEGG pathway, Hallmark, and WikiPathways) in the Molecular Signature Database (MSigDB v7.5.1) (39) using the fgsea package (v1.24) (40) in R (v4.2.2) with parameters of minimum size 15, maximum size 500, and 100,000 permutations. The statistical significance of the GSEA results was evaluated by adjusting the p-value using the BH procedure. GSEA results were visualized as a heatmap using pheatmap (v.1.0.12), and hierarchical clustering analysis of pathways was performed using Euclidean distance and the complete method (41).

## 3. Results

### 3.1 Physical and Chemical Properties of OA, HG, and GA

The physicochemical and molecular characteristics of OA, HG, and GA were compared, and molecular descriptors were calculated to quantitatively confirm the similarity of the compounds. According to Lipinski’s rule of five (42), OA and HG are druggable compounds except that they are hydrophobic, and GA is suitable for use as a drug under all conditions. In addition, OA and HG have similar chemical properties and the same chemical classification as pentacyclic triterpenoids; therefore, they are suitable for analysis as similar compounds (Table 1).

**Table 1.** Chemical properties of oleanolic acid, hederagenin, and gallic acid

|  | Chemical property | Property value | Druggability (Lipinski's rule) |
| --- | --- | --- | --- |
| Oleanolic acid | Molecular Weight | 456.7 | YES |
|  | XLogP3 | 7.5 | NO |
|  | Hydrogen Bond Donor Count | 2 | YES |
|  | Hydrogen Bond Acceptor Count | 3 | YES |
|  | Chemical class | Pentacyclic triterpenoid |  |
| Hederagenin | Molecular Weight | 472.7 | YES |
|  | XLogP3 | 6.8 | NO |
|  | Hydrogen Bond Donor Count | 3 | YES |
|  | Hydrogen Bond Acceptor Count | 4 | YES |
|  | Chemical class | Pentacyclic triterpenoid |  |
| Gallic acid | Molecular Weight | 170.12 | YES |
|  | XLogP3 | 0.7 | YES |
|  | Hydrogen Bond Donor Count | 4 | YES |
|  | Hydrogen Bond Acceptor Count | 5 | YES |
|  | Chemical class | Benzoic acids |  |

Lipinski's rule of five: Molecular Weight $\leq 500$ , XlogP3 $\leq 5$ , H-bond donors $\leq 5$ , H-bond acceptors $\leq 10$

Distance similarity calculations using molecular descriptors confirmed that OA and HG are similar to GA. The Euclidean distance analysis showed that the distance between OA and HG was approximately 40 times smaller than that between OA and GA and between HG and GA. In particular, as a result of calculating the cosine and Tanimoto distances, OA and HG were found to be more than 99% similar. Conversely, in the Tanimoto distance calculation results, GA had only approximately 7% similarity with the other two compounds, which quantitatively confirmed that it was a heterogeneous compound compared with the other two compounds (Table 2).

**Table 2.** Molecular descriptor similarity of the three components

|  | Euclidean distance | Cosine distance | Tanimoto distance |
| --- | --- | --- | --- |
| OA-HG | 4793.82 | 0.99995 | 0.99949 |
| OA-GA | 189942.02 | 0.76676 | 0.07616 |
| HG-GA | 194267.50 | 0.76446 | 0.07231 |

OA: Oleanolic acid; HG: Hederagenin; GA: Gallic acid

### 3.2 Results of Systems Pharmacology Analysis of OA, HG, and GA

Systems pharmacology analysis using the BATMAN-TCM database showed similar patterns for OA and HG, whereas GA showed different patterns. There were 44 druggable targets shared by OA and HG, and they shared interacting proteins at rates of 86% (OA:44/51) and 96% (HG:44/46). In contrast, only 20 % (5/25) of the GA protein was shared with the druggable targets of the other two compounds (Figure 2A, Supplementary Table 1).

**Figure 2.**
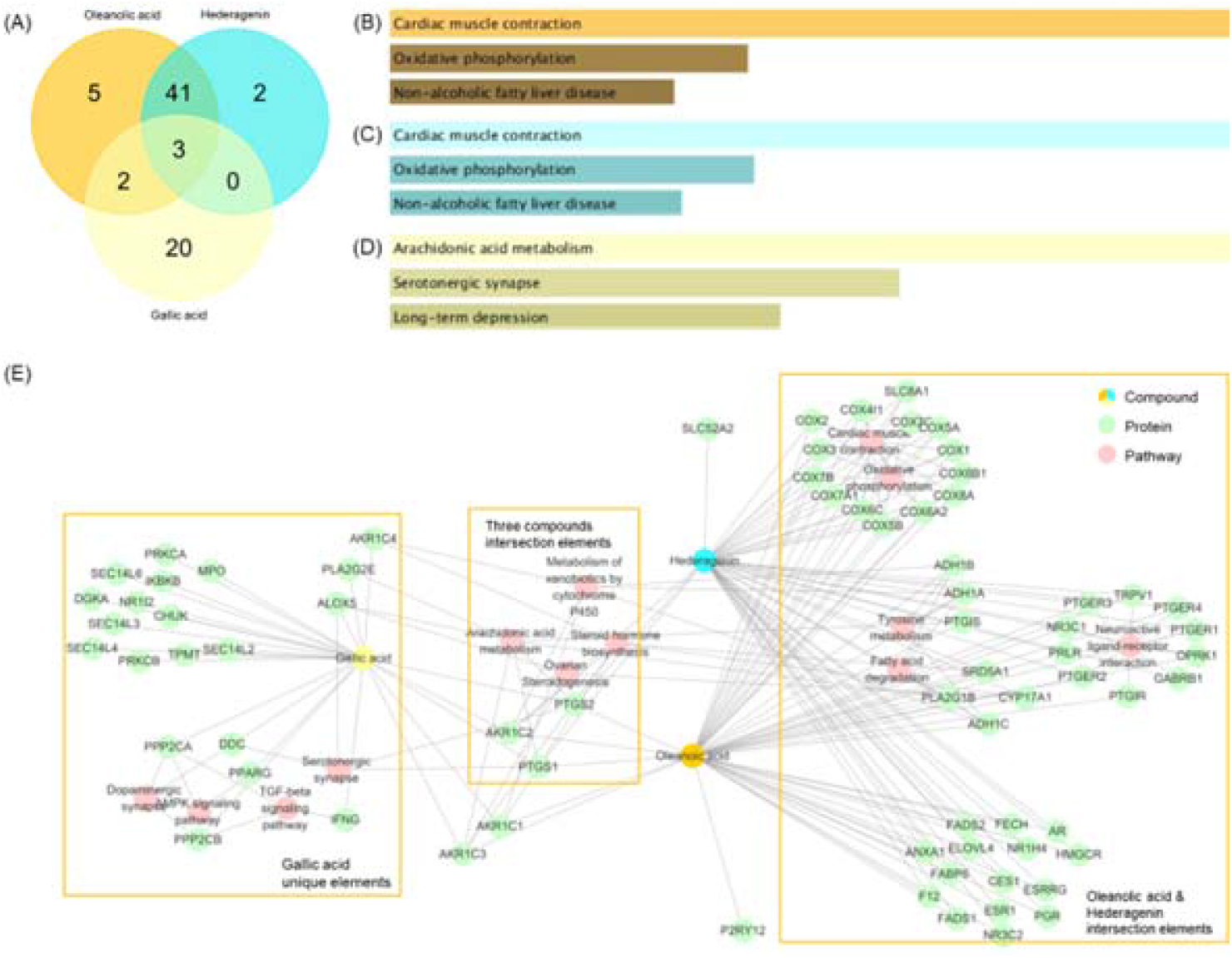
Systems pharmacology analysis of OA, HG and GA. A systems pharmacology platform, BATMAN-TCM, was used to predict druggable target proteins of oleanolic acid (OA), hederagenin (HG) and gallic acid (GA). Over-representation analysis (ORA) was performed on the EnrichR platform using the predicted druggable proteins and KEGG pathway gene sets, and a compound-protein-pathway (CPP) network was constructed using the ORA results. (A) Venn diagram shown using druggable target proteins of OA, HG, and GA predicted by BATMAN-TCM. (B) ORA results using KEGG pathway gene sets and druggable target proteins of OA. (C) ORA results using KEGG pathway gene sets and druggable target proteins of HG. (D) ORA results using KEGG pathway gene sets and druggable target proteins of GA. (E) CPP network constructed using OA, HG, GA target proteins and KEGG pathway analysis results. Orange, light blue, and yellow nodes represent OA, HG, and GA, respectively. Green nodes represent proteins interacting with compounds, and pink nodes represent valid pathways derived from proteins. The orange box indicates the classification of proteins and pathways, which are divided into 1) the action point where OA and HG act together, 2) the action point where GA acts alone, and 3) the action point where the three compounds act together.

In the ORA using the druggable target of each compound, the analysis results for OA and HG were consistent. The ORA of OA and HG showed that the three major pathways with the highest combined scores were cardiac muscle contraction, oxidative phosphorylation, and nonalcoholic fatty liver disease. In contrast, arachidonic acid metabolism, serotonergic synapses, and long-term depression were analyzed as major pathways in GA, and the results of the other two compounds were not consistent (Figure 2B-D). Even when the analysis results were expanded to the top 10 KEGG pathways, OA and HG matched nine pathways, whereas GA matched only the steroid hormone biosynthesis pathway with OA. In GO pathway analysis, both OA and HG were predicted to have the highest combined score for the mitochondrial electron transport mechanism. In addition, in the OMIM disease analysis, both OA and GA were predicted to act on cholesterol levels, migraine, and myocardial infarction (Supplementary Figure 1).

Network analysis results based on systems pharmacology platform analysis also showed that OA and HG share the same mechanism. OA and HG share the most druggable targets and mechanisms of cardiac muscle contraction, oxidative phosphorylation, neuroactive ligand-receptor interactions, tyrosine metabolism, and fatty acid degradation. However, GA had relatively many unique elements and, unlike the other two components, acted on serotonergic synapse, dopaminergic synapse, AMPK signaling pathway, and TGF-beta signaling pathway (Figure 2E).

### 3.3 Results of Molecular Docking Analysis of OA, HG and GA

Analysis of the top 50 druggable proteomes predicted by molecular docking analysis with OA, HG, GA, OA, and HG showed similar patterns (Supplementary Material 2). OA and HG shared 38 druggable proteins, while GA shared only 11 proteins with the other two (Figure 3A, Supplementary Table 2).

**Figure 3.**
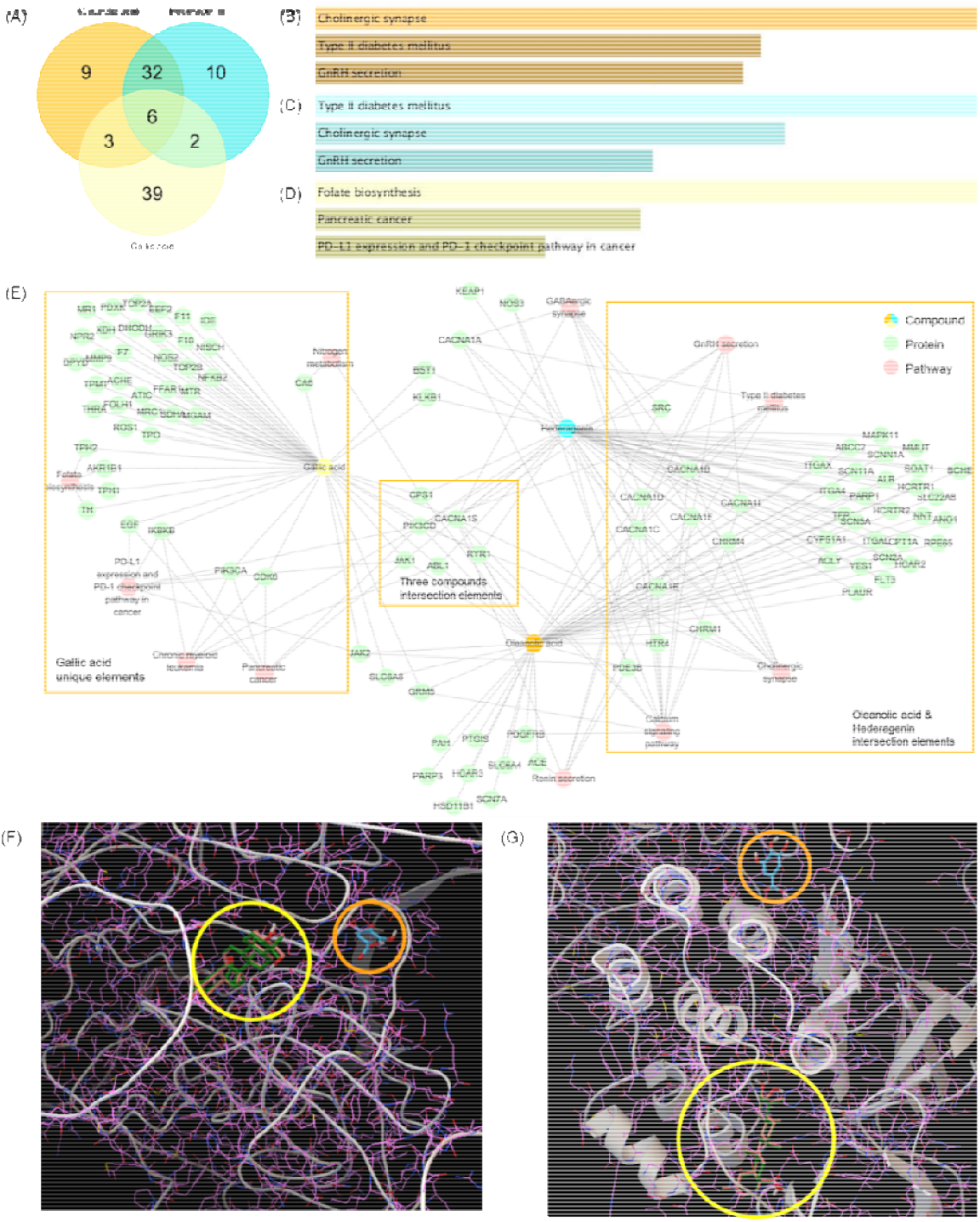
Results of molecular docking analysis of OA, HG and GA. Based on druggable proteome information, large-scale molecular docking analysis of oleanolic acid (OA), hederagenin (HG), and gallic acid (GA) was performed to predict compound-protein interactions. Over-representation analysis (ORA) was performed on the EnrichR platform using the predicted druggable proteins and KEGG pathway gene sets, and a compound-protein-pathway (CPP) network was constructed using the ORA results. Proteins interacting with all three compounds were additionally visualized as molecular docking analysis results. (A) Venn diagram of using 50 target proteins with the lowest binding affinity as a result of molecular docking analysis using OA, HG, and GA, respectively. (B) ORA results using KEGG pathway gene sets and druggable target proteins of OA. (C) ORA results using KEGG pathway gene sets and druggable target proteins of HG. (D) ORA results using KEGG pathway gene sets and druggable target proteins of GA. (E) CPP network constructed using OA, HG, GA target proteins and KEGG pathway analysis results. Orange, light blue, and yellow nodes represent OA, HG, and GA, respectively. Green nodes represent proteins interacting with compounds, and pink nodes represent valid pathways derived from proteins. The orange box indicates the classification of proteins and pathways, which are divided into 1) the action point where OA and HG act together, 2) the action point where GA acts alone, and 3) the action point where the three compounds act together. (F) Results of molecular docking analysis with ABL1 and three compounds. (G) Results of molecular docking analysis with JAK1 and three compounds. In the docking analysis, OA was shown in green, HG in pink, and GA in light blue. The yellow circle indicates the location where OA and HG interact, and the orange circle indicates the location where GA interacts.

In the ORA docking results, both OA and HG were predicted to act on cholinergic synapses, type 2 diabetes mellitus, and the gonadotropin-releasing hormone (GnRH) secretion pathway. In contrast, GA was predicted to act on pathways different from those of the two compounds, such as folate biosynthesis and the pancreatic cancer pathway (Figure 3B-D). Even when the analysis results were expanded to the top 10 KEGG pathways, OA and HG matched eight pathways, whereas GA did not match any pathway. In addition, in the GO pathway and OMIM disease analysis, both OA and HG showed consistent predictive results for membrane depolarization, calcium ion import, long QT syndrome, and hypertension. (Supplementary Figure 2).

In addition, network analysis based on molecular docking predicted that OA and HG share the same mechanism. Both OA and HG are predicted to act on GnRH secretion, type 2 diabetes mellitus, cholinergic synapses, and calcium signaling pathways. However, because GA had many proteins that interacted only with GA, unlike the other two compounds, it acted on pathways such as folate biosynthesis, pancreatic cancer, and chronic myeloid leukemia (Figure 3E).

Finally, the molecular docking analysis results of the proteins with which all three compounds interacted were visualized to confirm whether OA and HG bound to the same positions. Visualization confirmed that OA and HG interacted at similar positions in all proteins, whereas only GA interacted at different positions (Figure 3F, G, Supplementary Figure 3). These results suggest that OA and HG likely share similar mechanisms, and we can conclude that GAs may have different mechanisms even though they interact with the same protein.

### 3.4 Results of RNA-seq Analysis of OA, HG, and GA

By selecting the DEGs of OA, HG, GA, and a combination of OA and HG (COH) using RNA-seq analysis, it was confirmed that the expression patterns of OA, HG, and COH were similar (Supplementary Material 3). In particular, the 110 genes whose expression levels were altered by all three compounds were OA, HG, and COH. In contrast, among the 49 genes whose expression levels changed in GA, 43 showed a pattern of expression level change only in GA. (Figure 4A). Volcano plot analysis also confirmed that the expression levels of *FAM129A, PCK2, MTHFD2, PSAT1, ASNS, PHGDH, FGB*, and *EGLN3*, that were altered in COH, were altered in at least one of the OA and HG groups. However, it was not possible to identify genes whose expression levels changed in the same pattern as COH in the GA group (Figure 4B, Supplementary Figure 4).

**Figure 4.**
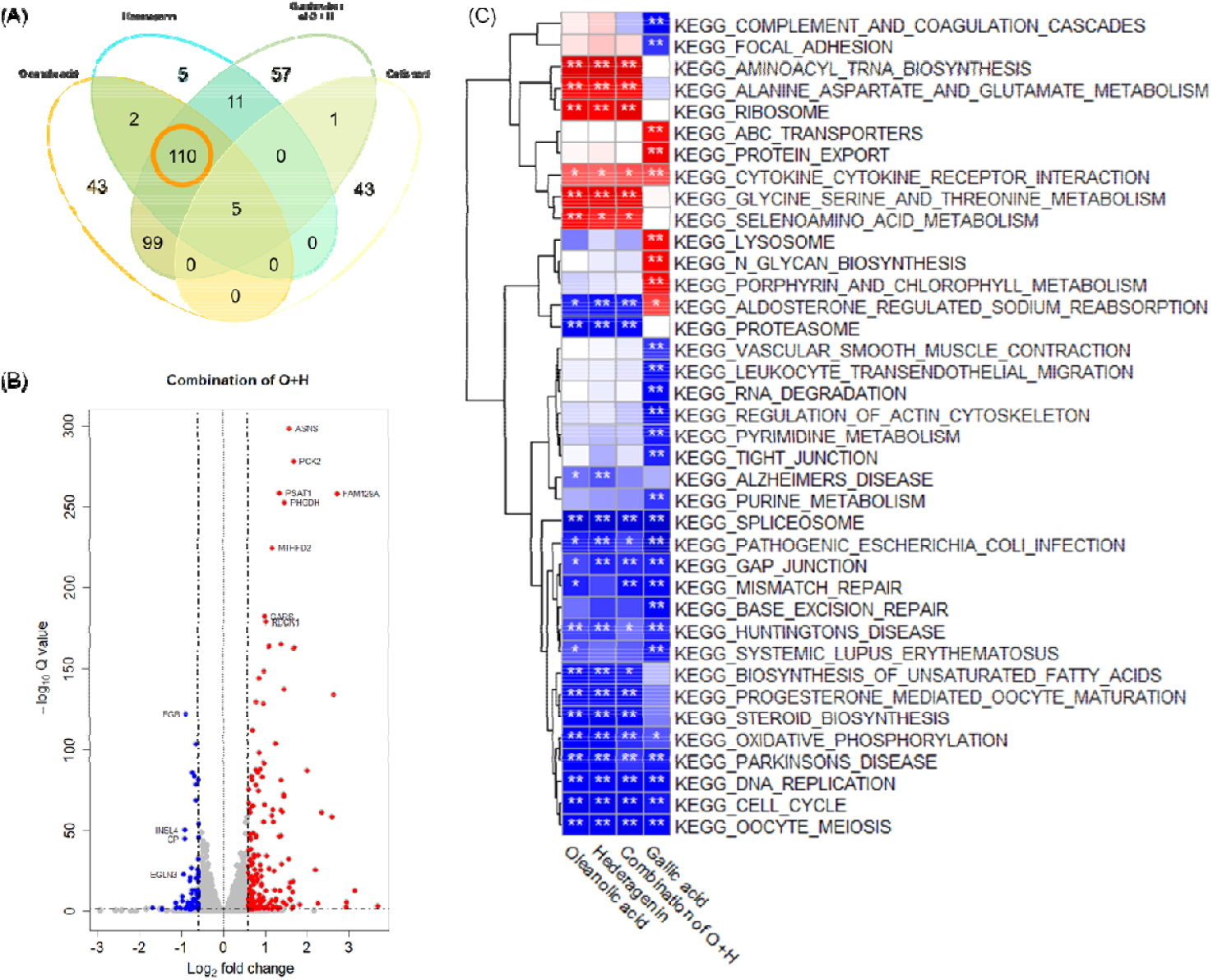
Results of RNA-seq analysis of OA, HG and GA. The A549 cell line was treated with oleanolic acid (OA), hederagenin (HG), a combination of OA and HG (COH), and gallic acid (GA), and the mechanism of action of each compound was derived using transcriptome data. Gene set enrichment analysis (GSEA) was performed using the gene expression results derived from each compound. (A) Venn diagram is shown using the genes selected as DEGs as a result of RNA-seq analysis of OA, HG, COH, and GA. Commonly selected DEGs in OA, HG, and COH are highlighted in orange circles. (B) A volcano plot was constructed using gene expression information derived from COH. (C) GSEA results of OA, HG, COH and GA were calculated using the KEGG pathway gene sets and RNA-seq analysis results of OA, HG, COH and GA.

In the GSEA results, OA, HG, and COH exhibited similar patterns. In KEGG pathway analysis, OA, HG, and COH showed pathways related to amino acid and fat metabolism, such as 1) Alanine, aspartate, and glutamate metabolism, 2) Glycine, serine, and threonine metabolism, 3) Biosynthesis of unsaturated fatty acids, and 4) Steroid biosynthesis. In contrast, GA was associated with RNA-related pathways, including 1) RNA degradation, 2) pyrimidine metabolism, and 3) purine metabolism (Fig 4C). In particular, OA, HG, and COH acted in opposition to GA in the aldosterone-regulated sodium reabsorption pathway.

## 4. Discussion and Conclusion

In this study, we confirmed that natural compounds with similar structures exhibit similar mechanisms of action. In other words, through analysis of the natural product-derived compounds OA, HG, and GA, it was found that OA and HG with similar structures had similar mechanisms of action, while GA and the other two compounds had different mechanisms of action.

The finding that compounds with similar structures exhibit similar mechanisms has three major novelties. First, an advanced network pharmacology research method was suggested by partially presenting a solution to overcome the difficulty of identifying multicomponent mechanisms, which are characteristic of natural products and an obstacle to new drug development. Second, a new natural product analysis method was proposed using large-scale molecular docking analysis. Finally, various biological data and analysis methods were used, such as in silico system pharmacology, docking analysis, and drug response RNA-seq. We confirmed the pharmacodynamic processes from the binding of natural compounds to druggable targets to the expression of drug efficacy through signaling by the bound using this method.

To date, most network pharmacology studies predicting the mechanisms of natural compounds have analyzed all components independently. However, as shown in the molecular descriptor similarity comparison, OA and HG were quantitatively similar compounds; therefore, they should not be considered independent components (Table 1,2). Rather, the two compounds should be considered dependent components that share the same mechanisms. Systematic pharmacological analyses confirmed that similar compounds share similar mechanisms, which was confirmed by molecular docking and transcriptome analyses. This result can reduce the complexity of mechanism prediction when analyzing multiple compounds. In other words, replacing similar compounds with the same scaffold as a representative compound is possible. This would help overcome the limitation posed to studying the mechanism by the large number of compounds in natural products. Natural products can effectively treat multifactorial diseases and comorbidities by acting on multiple targets. However, there is a disadvantage: it is difficult to use in developing new drugs because it is hard to predict the mechanism of natural products. If the complexity of mechanism prediction can be reduced using the method proposed in this study, it will greatly help natural-product-based drug development strategies. However, network pharmacological analysis showed that OA has a wider range of interacting targets than HG. Because the mechanism of action may vary depending on the differences in functional groups, polarity, and molecular weight, future research should be conducted to identify this.

Large-scale molecular docking analysis was performed using human-derived proteins. Because docking analyses based on the magic bullet paradigm are important for a small number of key targets, few attempts have been made to use a large number of human-derived protein structures. Natural products are not used after being subjected to preparing one active compound from a lead compound like in case of general medicines, but are instead used after being subjected to simple processing of plants, animals, and minerals. Therefore, among the many compounds present in natural products, it is important to confirm whether a certain compound interacts with a druggable protein. Because selecting a specific compound as a key compound in natural product mechanism research is difficult without specific criteria, a large-scale molecular docking analysis strategy is required. In particular, to study the treatment mechanisms of complex diseases by multiple compounds and targets of natural products, it is important to identify the target proteins that natural compounds bind to in the human body. When conducting such a mechanistic study, the large-scale docking analysis method used in this study will be important for future natural product research (Figure 3).

Finally, this study is significant in confirming all processes of a series of pharmacodynamic mechanisms of compounds in herbal medicine. It is well known that when compounds bind to target proteins in the human body, the second messenger is activated and affects the expression of new proteins. However, it is difficult to confirm the secondary drug activities of natural products using network pharmacology and molecular docking analyses. The secondary drug activity of the natural product, which the two analyses could not confirm, was confirmed using drug response transcriptome analysis. For example, the cholinergic synapse, GnRH secretion, and calcium signaling pathways obtained from the molecular docking analysis of OA and HG (Figure 3B, C) affect the aldosterone-regulated sodium reabsorption obtained from the RNA-seq analysis (Figure 4C). This is also consistent with the cardiac muscle contraction pathway obtained from the system pharmacology analysis. These results confirm that OA and HG affect heart health (Figure 2)(43). Biological data analysis shows a series of processes in which the human body absorbs natural compounds, interact with druggable targets, and is activated by secondary messengers. As shown in our study, the research method that analyzes systems biology, molecular docking, and even transcriptomes will be a milestone in analyzing the pharmacological mechanisms of natural products.

In this study, GSEA based on drug-response transcripts confirmed that similar compounds have similar mechanisms. A recent study identified the Aurora B pathway as a novel anticancer therapeutic activity of Paeoniae Radix (PR) extract in lung cancer cell lines via systematic transcriptome analysis (44). Among the eight compounds (HG, OA, GA, Albiflorin, Benzoic Acid, Catechin, Paeoniflorin, and Paeonol) in PR, HG and OA demonstrated anticancer efficacy by reducing Aurora kinase activity. In contrast, other compounds (GA, Benzoic Acid, Catechin, Paeoniflorin, and Paeonol) in PR induced cell-cycle arrest and apoptosis via p53 and MAPK activation. Although the structurally similar HG and OA were significantly altered by the Aurora B pathway, other compounds containing GA demonstrated anticancer effects through cell cycle arrest and apoptosis. The results of our study are consistent with previous findings showing that the structure of a compound determines its mechanism of action. These results demonstrate that the method used in this study can be effectively used to study the mechanisms of action of herbal medicines.

However, this study has several limitations. First, there was a limitation in not considering the number of compounds in the system pharmacology analysis. The dose of a drug is very important in all experiments, from *in vitro* experiments to clinical trials, because the biological mechanism and cell viability may vary depending on the dose. Systems pharmacology studies have been limited because they do not consider the doses of compounds contained in herbal medicines. The discovery in this study that compounds with the same scaffold have the same mechanism will have a positive impact on dose-based in silico herbal medicine research. However, this study only suggested a method for handling the dose of compounds in herbal medicines and did not consider the exact dose. In future studies, it will be necessary to propose a systematic pharmacological analysis strategy, such as LC-MS, that can quantitatively confirm the amounts of compounds.

Another limitation of our study is that the association between systems pharmacology, molecular docking, and RNA-Seq analysis was not clearly confirmed. All three methods have in common that they are analyzed to confirm the mechanism of action of the drug and analyzed using the same gene set. However, systems pharmacology and molecular docking are methods used to observe drug-target interaction mechanisms at the upstream level, whereas RNA-seq is a method used to observe drug-target interaction mechanisms at the downstream level after a second messenger system transmits a signal. Therefore, these methods maintain a consistent drug mechanism in some cases but are inconsistent in others because they involve complex interaction mechanisms. Therefore, further studies are required to connect and integrate different analytical methods based on the same compounds.

Nevertheless, the results of this study are meaningful in that they suggest an analysis strategy that can improve the existing systems pharmacology research analysis method using the result that natural product-derived compounds with the same scaffold have the same mechanism. In addition, this study is novel because it proposes a new natural product mechanism analysis method using large-scale molecular docking and confirms the results of secondary drug response transcripts using RNA-seq analysis.

## Acknowledgement

This research was funded by the research program of the Korea Institute of Oriental Medicine [grant number KSN1731122].

## Authorship contribution

M.S, P. contributed to the research design, analyzed data, and drafted the manuscript; S.J, B drafted the manuscript; S.M, P. analyzed data; J.M, Y conducted data design and production and drafted the manuscript; S.W, C. drafted and validated the manuscript. All authors have read and agreed to the published version of the manuscript.

## Data availability

The raw sequence and processed data were deposited in the NCBI Gene Expression Omnibus (GEO, https://www.ncbi.nlm.nih.gov/geo/) with accession number GSE228524.

