## supplementary figures and tables for "Comparative study of the mechanism of natural compounds with similar structures using docking and transcriptome data for improving in silico herbal medicine experimentations"

Supplementary Table 1 Prediction results of druggable target proteins of oleanolic acid, hederagenin, and gallic acid using the systems pharmacology platform BATMAN-TCM

| Names | #Total | Protein element |
| --- | --- | --- |
| OA & HG & GA | 3 | AKR1C2 PTGS1 PTGS2 |
| OA & HG | 41 | ADH1C ANXA1 AR CES1 COX1 COX2 COX3 COX4I1 COX5A COX5B COX6A2 COX6B1 COX6C COX7A1 COX7B COX7C COX8A CYP17A1 ELOVL4 ESR1 ESRRG F12 FABP6 FADS1 FADS2 FECH HMGCR NR1H4 NR3C1 NR3C2 OPRK1 PGR PLA2G1B PRLR PTGER1 PTGER2 PTGER3 PTGER4 SLC8A1 SRD5A1 TRPV1 |
| OA & GA | 2 | AKR1C1 AKR1C3 |
| OA | 5 | ADH1A ADH1B P2RY12 PTGIR PTGIS |
| HG | 2 | GABRB1 SLC52A2 |
| GA | 20 | AKR1C4 ALOX5 CHUK DDC DGKA IFNG IKBKB MPO NR1I2 PLA2G2E PPARG PPP2CA PPP2CB PRKCA PRKCB SEC14L2 SEC14L3 SEC14L4 SEC14L6 TPMT |
| OA: Oleanolic acid; HG: Hederagenin; GA: Gallic acid | | |

Supplementary Table 2 Prediction results of druggable target proteins of oleanolic acid, hederagenin, and gallic acid calculated using molecular docking analysis.

| Names | #Total | Protein element |
| --- | --- | --- |
| OA & HG & GA | 18 | ABL1 BST1 CACNA1S CACNA2D3 CPS1 DPYD GRM5 JAK1 KEAP1 NOS2 PARP3 PIK3CD RPE65 RYR1 SCN7A SLC6A8 TOP2B VWF |
| OA & HG | 65 | ABCA1 ABCC2 ACE ACLY ALB AMY2A ANO1 AR ATP4A BCHE BRAF C5 CACNA1B CACNA1C CACNA1D CACNA1E CACNA1F CACNA1H CACNA1I CCR4 CHRM1 CHRM4 CHRM5 CPT1A CSF1R CSF3R DNMT1 FLT3 GAA GLP2R GRIN2B HCAR2 HCRTR1 HCRTR2 HSD11B1 HTR4 ITGA4 ITGAL ITGAX JAK3 KCNMA1 MAPK11 MMUT NNT NOS1 NOS3 PDE3B PDGFRB PLAUR PTGIS SCN11A SCN2A SCN4A SCN5A SCN9A SCNN1A SLC22A8 SLC38A4 SLC6A4 SOAT1 SRC TFRC TLR9 TRPV3 YES1 |
| OA & GA | 4 | JAK2 MGAM MRC1 PAH |
| HG & GA | 5 | CACNA1A KLKB1 NOD2 PARP1 RAF1 |
| OA | 13 | C4B CACNA1G CACNG1 DRD2 ERBB2 FGFR1 FRK GCGR HCAR3 RARB SCN10A SLC22A6 SLC52A2 |
| HG | 12 | ADORA1 ATP2B3 CACNA2D4 CYP51A1 DHFR ERBB4 GRIN2A GSR KCNH6 NTRK3 PDE4A SCNN1D |
| GA | 73 | ACHE ADRB2 AKR1B1 AKR1C2 AKR1D1 ALOX5 ATIC ATP2A2 BCR C3 CA6 CACNA2D1 CACNB2 CCND1 CDK6 DHODH EEF2 EGF F10 F11 F7 FDPS FFAR1 FOLH1 GRIA1 GRIK3 HRH1 HSD17B10 HTR7 IDE IKBKB IMPDH2 ITGA2B MAP3K1 MAPK1 MME MMP9 MR1 MTR NFKB2 NISCH NPR1 NPR2 NR3C1 OPRM1 PCSK9 PDXK PIK3CA PIK3CG PLAT PNP PRKAA1 PRKCE RET ROS1 RYR2 SDHA SLC38A2 SLC5A2 SLC7A7 SLC8A1 TH THRA TOP2A TPH1 TPH2 TPMT TPO TRPM8 TYMP TYR UGCG XDH |
| OA: Oleanolic acid; HG: Hederagenin; GA: Gallic acid | | |

Supplementary Table 3 Differentially expressed gene selection for oleanolic acid, hederagenin, and gallic acid using RNA-seq analysis

| Names | #Total | Protein element (RNA-seq DEG) |
| --- | --- | --- |
| OA & HG &  Combi & GA | 5 | GREM1 KIF1A MARCHF4 RP11-1212A22.4 SLC26A10 |
| OA & HG &  Combi | 110 | AARS ABCG1 ADM2 AKNA ALDH1L2 ALDOC ANK2 AP1M2 APLF APOL6 ARGEF2 ASNS ATF3 AXIN2 BCAT1 BEX2 BTC CA9 CAPN5 CARS CBS CCND3 CDH4 CEBPG CHAC1 CLGN CLIC3 CSF2RA CSTA CYP4V2 DDIT3 DDIT4 EGLN3 EIF4EBP1 EPHA7 FAM129A FBXL16 FLRT1 FUT1 GARS GDF15 GPT2 HOXB9 HRK HYAL1 IARS ICAM1 IFRD1 IL11 INHBE INSL4 IRAK2 JDP2 JUN KCNH1 KIF21B KLB KLHL3 LAMC2 LAMP3 LY6K MARS MCTP1 MTHFD2 NDUFA4L2 NIPAL4 NUPR1 OGT PABPC1L PCK2 PCSK9 PHGDH PIP5KL1 PLA2G4C PMAIP1 PRKG2 PRTN3 PSAT1 PSMA1 PYCR1 RAB39B RBCK1 RGPD6 ROBO3 SARS SERPINE1 SESN2 SLC1A4 SLC43A1 SLC5A5 SLC6A9 SLC7A1 SLFNL1 SMOC1 SNTB1 SORCS2 SPX STC2 TM6SF1 TRIB3 TSC22D3 TUBB4A TUBE1 ULBP1 UNC5B VLDLR XPOT YARS ZNF674 ZNF853 |
| OA & Combi | 99 | ABLIM3 ADAT3 AJUBA ANKRD1 ANXA3 APBB2 APOL1 ARRB1 ASGR1 ASS1 ATF4 BBC3 BCAS1 C2orf82 C6orf48 C9orf91 CABYR CALR CCDC113 CCT6B CDC42EP1 CDCP1 CDKN2D CITED4 CP CPEB1 CTPS1 CXCL2 CXCL5 CXCL8 DMRTA1 DOC2B DPEP1 DPY19L2 EREG ERN1 ESAM FADS2 FAM132B FAM228B FAM83A FAM86B1 FGA FGB FGG FOSL1 FYN GADD45A GAL3ST1 GFPT2 GPD1L GPR27 GPX2 GRB10 HMGN5 HSPA1A HTR2B IL20RB KCNG1 KDM7A KIAA1161 KRT80 LAMA4 MGAM MOCOS MORN4 MT-ND6 N4BP3 NAP1L2 NCCRP1 NDRG2 NEB NECAB2 NES NFKB2 NGEF NPAS1 NSUN7 OPLAH OSBPL6 PALM3 PEAR1 PLAU PPFIA4 PPP1R15A PYGB SH2D2A SLC16A3 SLC29A4 STK33 SYTL1 TCEA1 TF TMEM79 TNFRSF9 TUBB4B VAV3 WARS XDH |
| HG & Combi | 11 | AGMO C1orf61 CIDEB COL16A1 LONRF3 NR4A1 RP11-644F5.10 RP11-849F2.7 RP11-903H12.5 RPS6KA2 SERPINA3 |
| GA & Combi | 1 | C4B |
| OA & HG | 2 | KLHL13 UGT1A6 |
| OA | 43 | ACYP1 ASB4 BAAT BEX4 BIN2 C2orf88 CDRT4 CHRFAM7A DACT2 DRD1 FAM166B HHIP HPDL HSPA13 HSPA1B KLF11 KLHL35 LCN2 MPP2 NALCN NMRK1 NPIPA7 OSBP2 P2RY6 PDGFB PHOSPHO1 PODXL PRSS35 RASA4B RBFOX3 RHOV RP11-468E2.2 SERPINB8 SLC16A14 SP140L TAS2R20 TGFA THBD TRIM73 TSEN15 TUBB2B VDR ZNF608 |
| HG | 5 | GOLT1A PRDM13 RP11-315D16.2 ST20-MTHFS ZNF239 |
| Combination of OA & HG | 57 | ALOX12B ARMCX4 CARF CCDC146 CDKN2C CSNK2B-LY6G5B-562 DFNB59 DHCR24 DNAJB5 DUSP8 DYSF EMP2 F7 FAM111B FBLN5 FHL2 FJX1 GOLGA8O GOT1 HERPUD1 HIC1 HLA-DMA IGDCC4 IL15RA KLHDC7B LONP1 LOXL2 LRG1 MICB MSC MT-CO3 MT-ND5 MUC16 NAV2 NCALD NKX2-8 NR4A3 OAS1 ODF3B PCDHGC3 RAB3A RASL11A RELB RNF43 RP11-1220K2.2 RP11-159D12.5 RP11-176H8.1 RP11-407N17.3 SAMD4A SLC1A5 SLC6A15 SMOX SYTL5 TNFRSF12A TTLL11 VAV1 VEGFA |
| GA | 43 | ACOT4 ADPRH AGMAT ALPK3 ANPEP ARHGEF5 CAND2 CASKIN1 CDK5R2 CHRNA3 CSPG5 CYB5R2 DHRS2 DNAJB9 DTX3 EMILIN2 EML6 EPCAM EPPK1 FLNC GIPR GLUL HENMT1 HMOX1 MAGI2 MAP6 MARCHF9 PPP1R3G PRODH2 PRTFDC1 RAPGEF3 RYR1 SALL4 SEC24D SLC1A1 SLC22A31 SLX1A SULT1A3 SYT7 TMEM98 TTC39A ZNF469 ZNF544 |
| OA: Oleanolic acid; HG: Hederagenin; Combi: Combination of Oleanolic acid and Hederagenin; GA: Gallic acid | | |

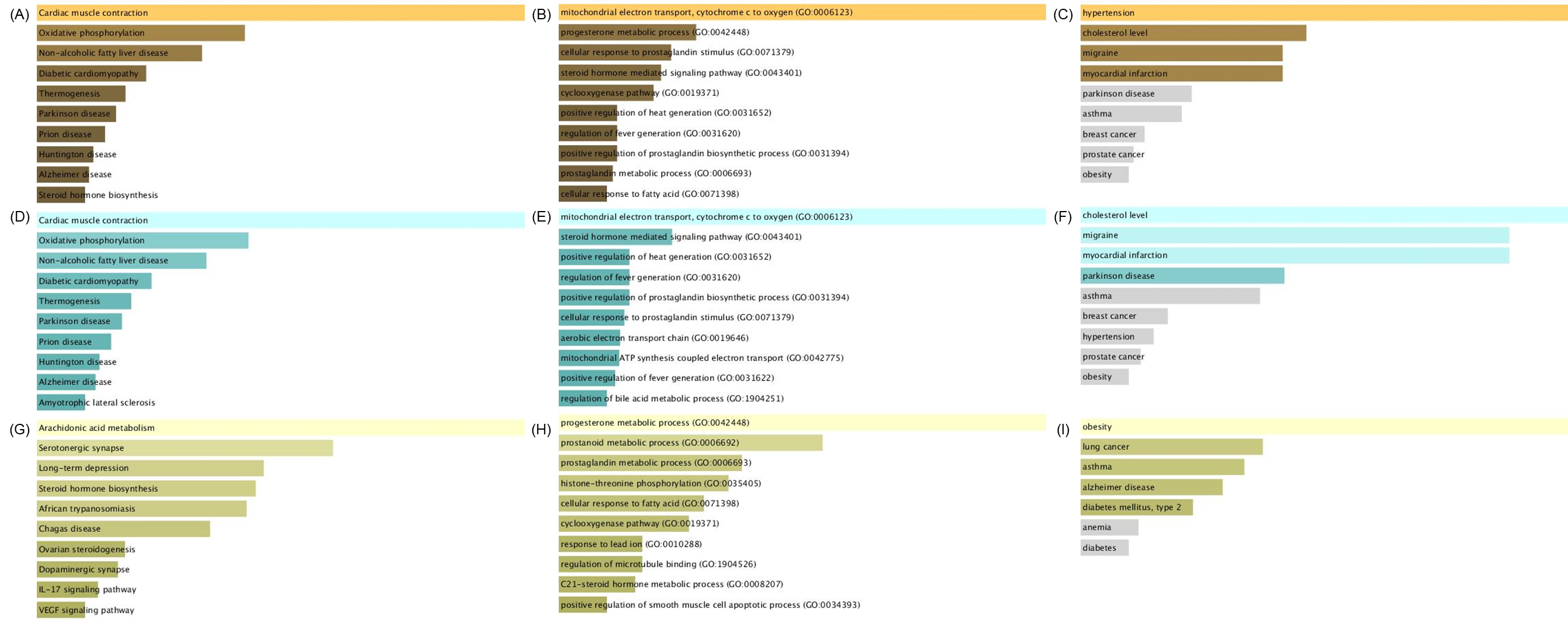

**Supplementary Figure 1 Over-representation analysis (ORA) using systems pharmacology platform.** Results of ORA on the EnrichR platform using druggable proteins of oleanolic acid (OA), hedergenin (HG) and gallic acid (GA) predicted using the systems pharmacology platform BATMAN-TCM. Only the top 10 pathways with the highest combined score were shown. (A) ORA results using KEGG pathway gene sets and druggable target proteins of OA. (B) ORA results using GO biological process gene sets and druggable target proteins of OA. (C) ORA results using OMIM disease gene sets and druggable target proteins of OA. (D) ORA results using KEGG pathway gene sets and druggable target proteins of HG. (E) ORA results using GO biological process gene sets and druggable target proteins of HG. (F) ORA results using OMIM disease gene sets and druggable target proteins of HG. (G) ORA results using KEGG pathway gene sets and druggable target proteins of GA. (H) ORA results using GO biological process gene sets and druggable target proteins of GA. (I) ORA results using OMIM disease gene sets and druggable target proteins of GA.

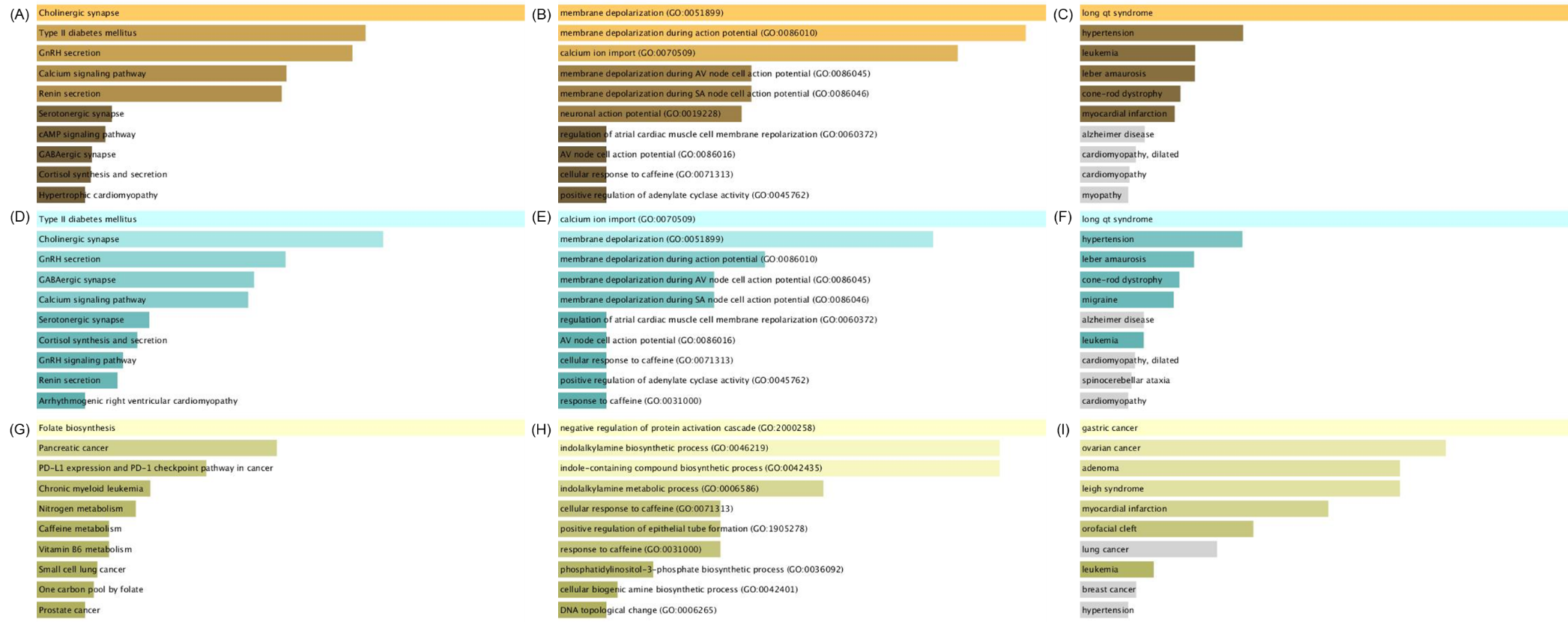

**Supplementary Figure 2 Results of over-representation analysis (ORA) using molecular docking.** These are the results of performing ORA on the EnrichR platform using 50 proteins predicted by molecular docking analysis with the lowest binding affinity to oleanolic acid (OA), hedergenin (HG), and gallic acid (GA). Only the top 10 pathways with the highest combined score were shown. (A) ORA results using KEGG pathway gene sets and druggable target proteins of OA. (B) ORA results using GO biological process gene sets and druggable target proteins of OA. (C) ORA results using OMIM disease gene sets and druggable target proteins of OA. (D) ORA results using KEGG pathway gene sets and druggable target proteins of HG. (E) ORA results using GO biological process gene sets and druggable target proteins of HG. (F) ORA results using OMIM disease gene sets and druggable target proteins of HG. (G) ORA results using KEGG pathway gene sets and druggable target proteins of GA. (H) ORA results using GO biological process gene sets and druggable target proteins of GA. (I) ORA results using OMIM disease gene sets and druggable target proteins of GA.
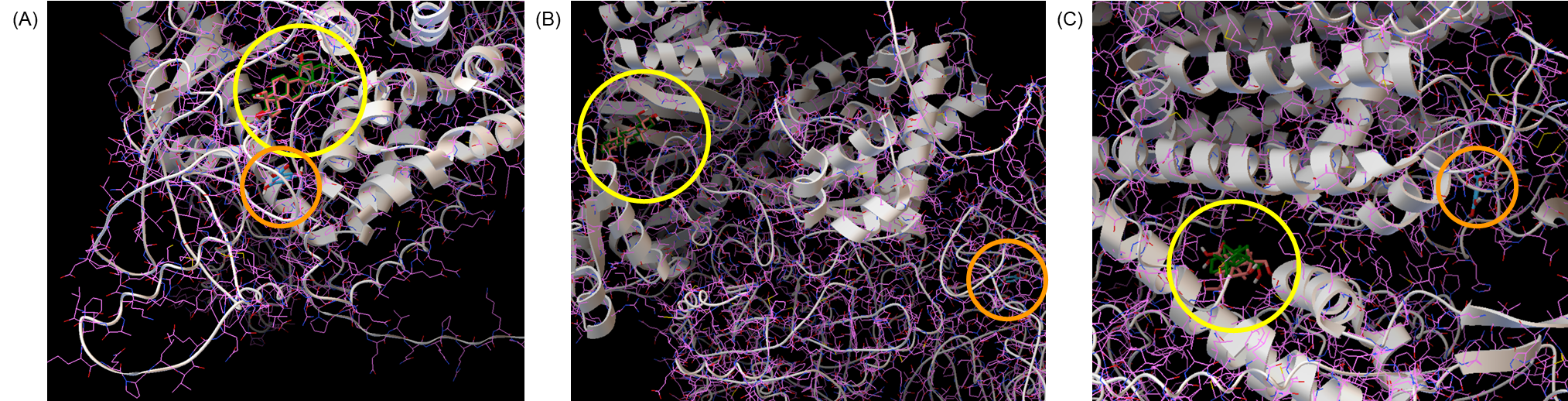

**Supplementary Figure 3 Results of molecular docking analysis and visualization.** Proteins interacting with all three compounds were visualized as molecular docking analysis results. (A) Results of molecular docking analysis with PIK3CD and Oleanolic acid (OA), Hederagenin (HG), and Gallic acid (GA). (B) Results of molecular docking analysis with CPS1 and OA, HG, and GA. (C) Results of molecular docking analysis with CACNA1S and OA, HG, and GA. OA was shown in green, HG in pink, and GA in light blue. The yellow circle indicates the location where OA and HG interact, and the orange circle indicates the location where GA interacts.

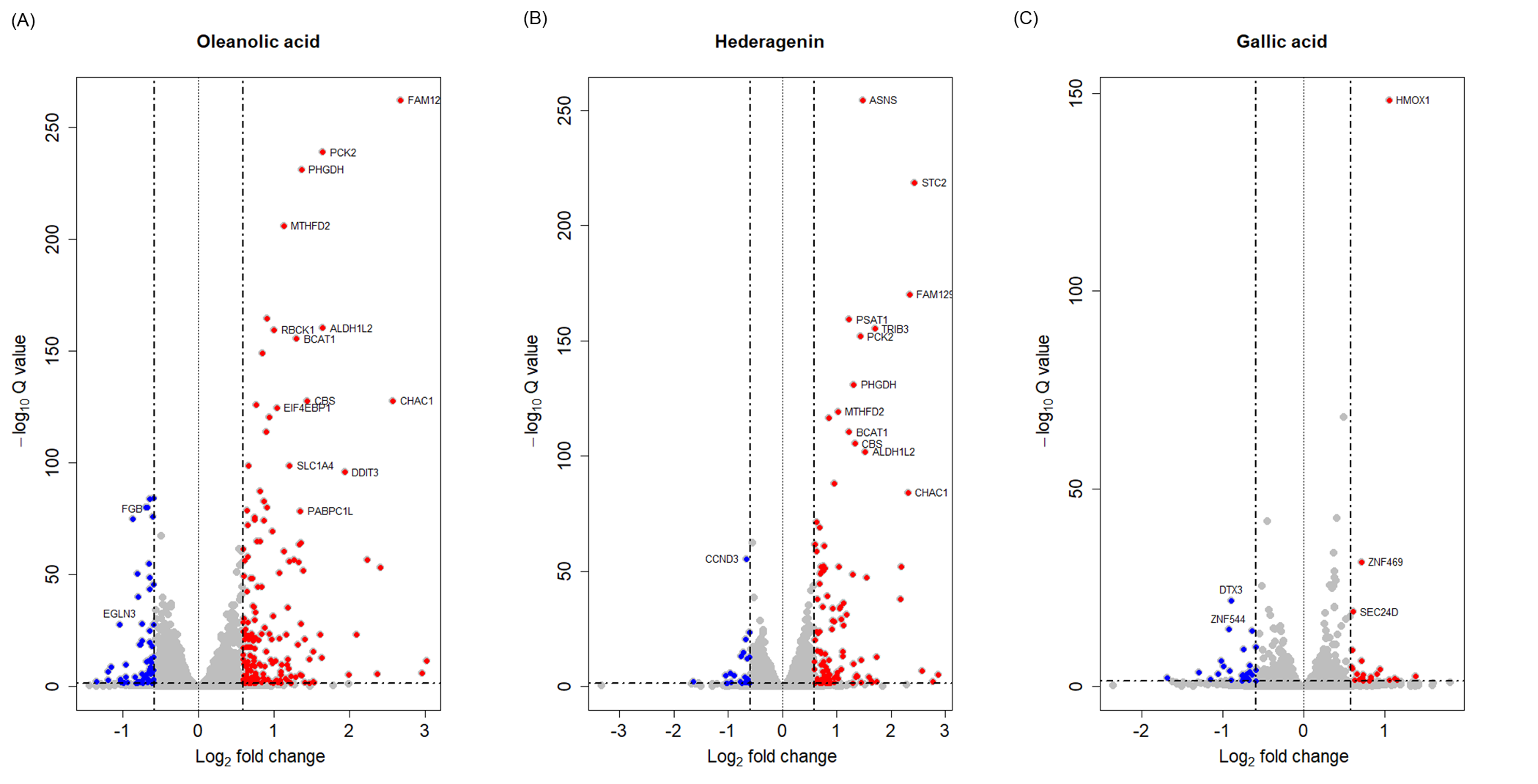

**Supplementary Figure 4 Volcano plots constructed using differentially expressed genes (DEGs) results.** The A549 cell line was treated with oleanolic acid (OA), hederagenin (HG), and gallic acid (GA), and DEGs of each compound were derived using transcriptome data. Volcano plots were constructed using genes selected as DEGs in OA, HG and GA, respectively. (A) A volcano plot was constructed using DEGs derived from OA. (B) A volcano plot was constructed using DEGs derived from HG. (C) A volcano plot was constructed using DEGs derived from GA.

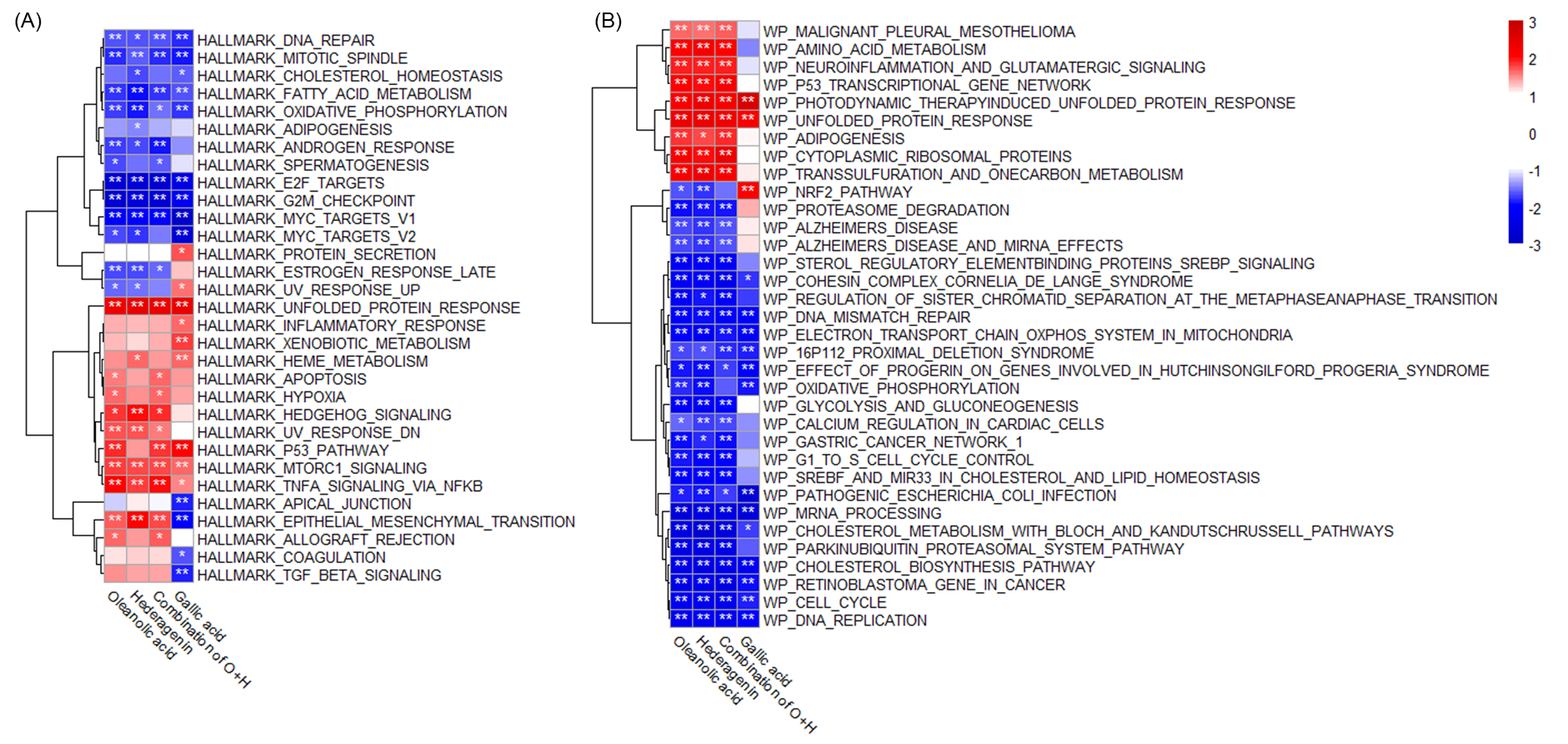

**Supplementary Figure 5 Result of gene set enrichment analysis (GSEA) performed using gene expression.** The A549 cell line was treated with oleanolic acid (OA), hederagenin (HG), a combination of OA and HG (COH), and gallic acid (GA), and gene expression information of each compound was confirmed using transcriptome data. Gene set enrichment analysis (GSEA) was performed using the gene expression results derived from each compound. (A) GSEA results of OA, HG, COH and GA were calculated using the Hallmark gene sets and RNA-seq analysis results of OA, HG, COH and GA. (A) GSEA results of OA, HG, COH and GA were calculated using the Wikipathway gene sets and RNA-seq analysis results of OA, HG, COH and GA.
